# Tensor-cell2cell v2 unravels coordinated dynamics of protein- and metabolite-mediated cell-cell communication

**DOI:** 10.1101/2022.11.02.514917

**Authors:** Erick Armingol, Reid O. Larsen, Lia Gale, Martin Cequeira, Hratch Baghdassarian, Nathan E. Lewis

**Affiliations:** Bioinformatics and Systems Biology Graduate Program, University of California San Diego, La Jolla, CA 92093, USA; Biomedical Sciences Graduate Program, University of California San Diego, La Jolla, USA; Department of Pharmacology, University of California San Diego, La Jolla, USA; Center for Molecular Medicine, Complex Carbohydrate Research Center, and Department of Biochemistry and Molecular Biology, University of Georgia, Athens, USA; Department of Pediatrics, University of California San Diego, La Jolla, USA; Department of Bioengineering, University of California San Diego, La Jolla, USA

## Abstract

**Summary:** Cell-cell communication dynamically changes across time while involving diverse cell populations and ligand types such as proteins and metabolites. While single-cell transcriptomics enables its inference, existing tools typically analyze ligand types separately and overlook their coordinated activity. Here, we present Tensor-cell2cell v2, a computational tool that can jointly analyze protein- and metabolite-mediated communication over time using coupled tensor component analysis, while preserving each modality of inferred communication scores independently, as well as their data structures and distributions. Applied to brain organoid development, Tensor-cell2cell v2 uncovers dynamic, coordinated communication programs involving key proteins and metabolites across relevant cell types across specific time points.

**Availability and implementation:** Tensor-cell2cell v2 and its new coupled tensor component analysis are implemented in Python and available as part of the cell2cell framework at https://github.com/earmingol/cell2cell. This python library is available on PyPI. Analyses of this manuscript can be reproduced in a Code Ocean capsule at https://doi.org/10.24433/CO.0061424.v1 and online tutorials can be found at https://cell2cell.readthedocs.io.

**Supplementary information:** Supplementary data are available at *bioRxiv* online.

## 1 Introduction

Cell-cell communication (CCC) is multimodal and dynamic, involving both small molecule and protein signals that act with fine spatiotemporal coordination to drive multicellular functions (Armingol *et al*., 2021). Although single-cell transcriptomics has enabled computational methods to infer cell-cell communication from gene expression (Almet *et al*., 2021; Armingol *et al*., 2024; Cesaro *et al*., 2025), most of them are focused on protein-protein interactions defining ligand-receptor (LR) pairs that cells can use to communicate.

Recent efforts have helped to incorporate the use of metabolite ligands for inferring CCC from single-cell transcriptomics (Zhang *et al*., 2024; Dimitrov *et al*., 2024; Troulé *et al*., 2025; Zheng *et al*., 2025). However, they do not account for both protein- and metabolite-mediated CCC simultaneously, often evaluating them separately in downstream analyses. Protein-mediated CCC can be directly inferred from gene expression, whereas metabolite-mediated CCC is inferred indirectly from enzymes that produce or consume the small molecules (Zheng *et al*., 2025). Consequently, protein- and metabolite-based scores are only informative in isolation as they differ in nature and scale, limiting their joint interpretation. Moreover, existing tools do not consider temporal coordination of LR pairs, preventing insights of protein and metabolite ligands acting in concert.

Coupled factorization approaches that link either tensors or matrices have demonstrated utility in revealing coordinated patterns across multimodal biological datasets while preserving the inherent data structure of each modality (Tan and Meyer, 2024). For example, such approaches have been used to jointly analyze static and dynamic metabolomics data, revealing shared subject patterns linked to biomarker profiles from both modalities (Li *et al*., 2024). To find dynamic CCC patterns driven by biological contexts, we previously introduced Tensor-cell2cell, a tool that applies tensor component analysis (TCA) on one modality of CCC and enables the incorporation of time as the biological context (Armingol, Baghdassarian, *et al*., 2022). Expanding this idea, coupled tensor factorization is particularly suited for studying multimodal CCC, while also considering shared changes across modalities in relation to cellular contexts (e.g., time points). Thus, coupled tensor component analysis (CTCA) provides a powerful strategy to integrate protein- and metabolite-mediated signals, capturing shared dynamics while preserving the distinct score distributions and biological insights of each modality.

Here, we present Tensor-cell2cell v2, a new version of our tool enabling CTCA to perform a joint evaluation of protein and metabolite ligand-receptor activity across time while maintaining each modality’s data structure and score distribution. We applied it to cortical brain organoid data, uncovering time-driven communication programs coordinating both ligand types during cortical development.

## 2 Methods

### 2.1 Preprocessing of the RNA-seq data

We analyzed single-cell RNA-seq data from cortical brain organoids at 1, 3, 6, and 10 months (Trujillo *et al*., 2019) available under GEO accession number GSE130238. To reconstruct the original study annotations based on pertinent markers, we processed this data with Seurat v3.0 (Stuart et al., 2019). Genes detected in fewer than 3 cells per time point were removed. After merging time points, we retained cells with 200-6000 genes and <7.5% mitochondrial content. Data were normalized, log-transformed, and scaled. Principal component analysis (PCA) was performed from highly variable genes identified with the FindVariableFeatures function. Batch correction was done using harmony with default parameters (Korsunsky et al., 2019). Cells were clustered using the top 20 harmony components (resolution = 0.5), yielding 17 clusters, which we merged to reconstitute the 7 cell types in the original study: glutamatergic, GABAergic, glia, progenitor, intermediate progenitor (IP), mitotic, and other cells (Supplementary Figure 1). Similar to that work, mitotic and other cells were excluded from further analysis.

### 2.2 Inferring cell-cell communication

Gene expression raw counts were normalized by library size and log-transformed to obtain log1p(CP10k) values. These were then used as input for cell2cell (Armingol, Ghaddar, *et al*., 2022) and MEBOCOST (Zheng *et al*., 2025) to infer protein- and metabolite-mediated CCC, respectively. Both tools compute communication scores based on LR interactions: for protein-mediated CCC, we used CellChatDB (Jin *et al*., 2021), which contains 2,005 human protein-based LR pairs; for metabolite-mediated CCC, we used the MEBOCOST database (Zheng *et al*., 2025), which includes 782 metabolite-based ligand-sensor interactions. For each modality, we constructed a 4D communication tensor from the resulting scores, organized by time points, LR pairs, and sender and receiver cell types (Armingol, Baghdassarian, *et al*., 2022).

### 2.3 Non-negative coupled tensor component analysis

Building upon a non-negative TCA (Williams *et al*., 2018), we implemented a non-negative CTCA to simultaneously factorize two non-negative tensors sharing all but one mode (i.e., dimension). To detail this approach, let *χ* and *χ ′* represent two tensors of size *C* x *P* x *S* x *T* and *C* x *P’* x *S* x *T*, respectively, where *C, S* and *T* correspond to the shared number of contexts/samples, sender cells and receiver cells, while *P* and *P’* correspond to the tensor-specific number of ligand-receptor pairs (or any other private dimension), representing protein- and metabolite-based LR pairs, respectively. Hence, *χ* _*ijkl*_ and *χ ′*_*imkl*_ denote the representative CCC in context *i*, using the protein- and metabolite-based LR pairs *j* and *m* respectively, between the sender cell *k* and receiver cell *l*.

The CTCA method corresponds to a coupled version of the CANDECOMP/PARAFAC decomposition (Carroll and Chang, 1970; Harshman and Others, 1970), yielding the simultaneous factorization of *χ* and *χ ′*, each through a sum of *R* tensors of rank-1:

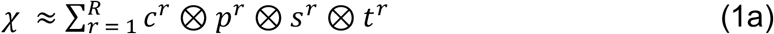

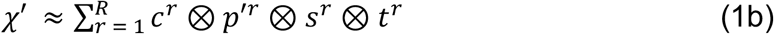

Where the notation ⊗represents the outer product and *c*^*r*^, *s*^*r*^*and t*^*r*^are shared vectors across both tensors, while *p*^*r*^ and *p′*^*r*^ are tensor-specific vectors for the private dimension (here, LR pairs). All vectors contain non-negative loadings representing the importance of each element in the corresponding factor *r*. Values of individual elements in these approximations are represented by:

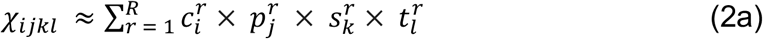

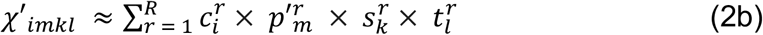

The coupled tensor factorization is performed by iterating the following objective function until convergence through an alternating least squares minimization (Anandkumar *et al*., 2014; Rabanser *et al*., 2017):

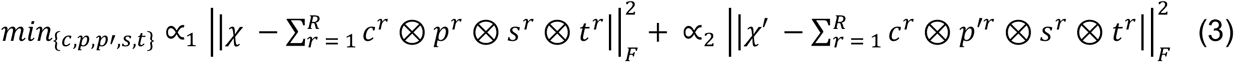

Where 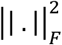 represents the squared Frobenius norm, and ∝ and ∝ are balancing weights. These weights are computed to compensate for size differences in the private dimension. Specifically, if *N*_1_ and *N*_2_ are the sizes of the private dimension in each tensor (number of protein- and metabolite-receptor interactions, respectively), we set *w*_1_ = (*N*_1_ + *N*_2_)/*N*_1_ and *w*_2_ = (*N*_1_ + *N*_2_)/*N*_2_. Then, ∝_1_= *w*_1_/(*w*_1_ + *w*_2_) and ∝_2_= *w*_2_/(*w*_1_ + *w*_2_), representing balancing weights that sum to one. This inverse weighting ensures that tensors with fewer elements in the private dimension receive proportionally higher weight, preventing larger tensors from dominating the optimization. This is particularly important when comparing modalities with vastly different numbers of elements (e.g., more protein than metabolite LR pairs, or vice versa), ensuring both conditions contribute equally to the discovered patterns.

### 2.4 Measuring the error of the CTCA

To quantify the error of the CTCA in representing the original 4D communication tensors (*χ* and *χ ′*) through the lower rank tensors (sum of *R* tensors of rank-1 in equations 1a,b), we use a combined normalized reconstruction error. First, individual normalized errors are computed for each tensor:

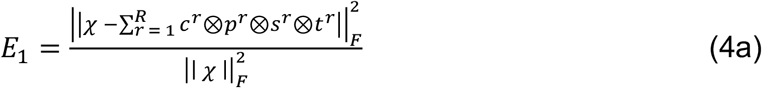

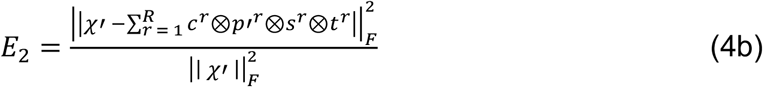

Then, the combined error *E* is calculated as a weighted average:

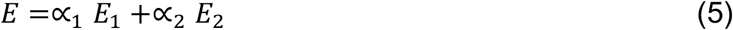

Where ∝_1_ and ∝_2_ represent the balancing weights in equation (3). This ensures both tensors contribute equally to the combined error metric. This combined error maintains the interpretability of the standard normalized error, acting analogously to the fraction of unexplained variance (Williams *et al*., 2018).

## 3 Results

We extended the functionalities of Tensor-cell2cell, a TCA framework for identifying context-driven communication programs, to investigate how CCC mediated by proteins and metabolites is coordinated. Briefly, Tensor-cell2cell summarizes each communication pattern through latent factors that capture combinations of sender-receiver cell pairs and their signaling molecules, while linking them to specific dynamics across biological contexts. In this second version, we implemented a non-negative CTCA devised to jointly analyze protein- and metabolite-mediated CCC across time (Figures 1A-C), overcoming the limitations of existing approaches that analyze these modalities separately and do not account for a temporal dimension.

**Figure 1.**
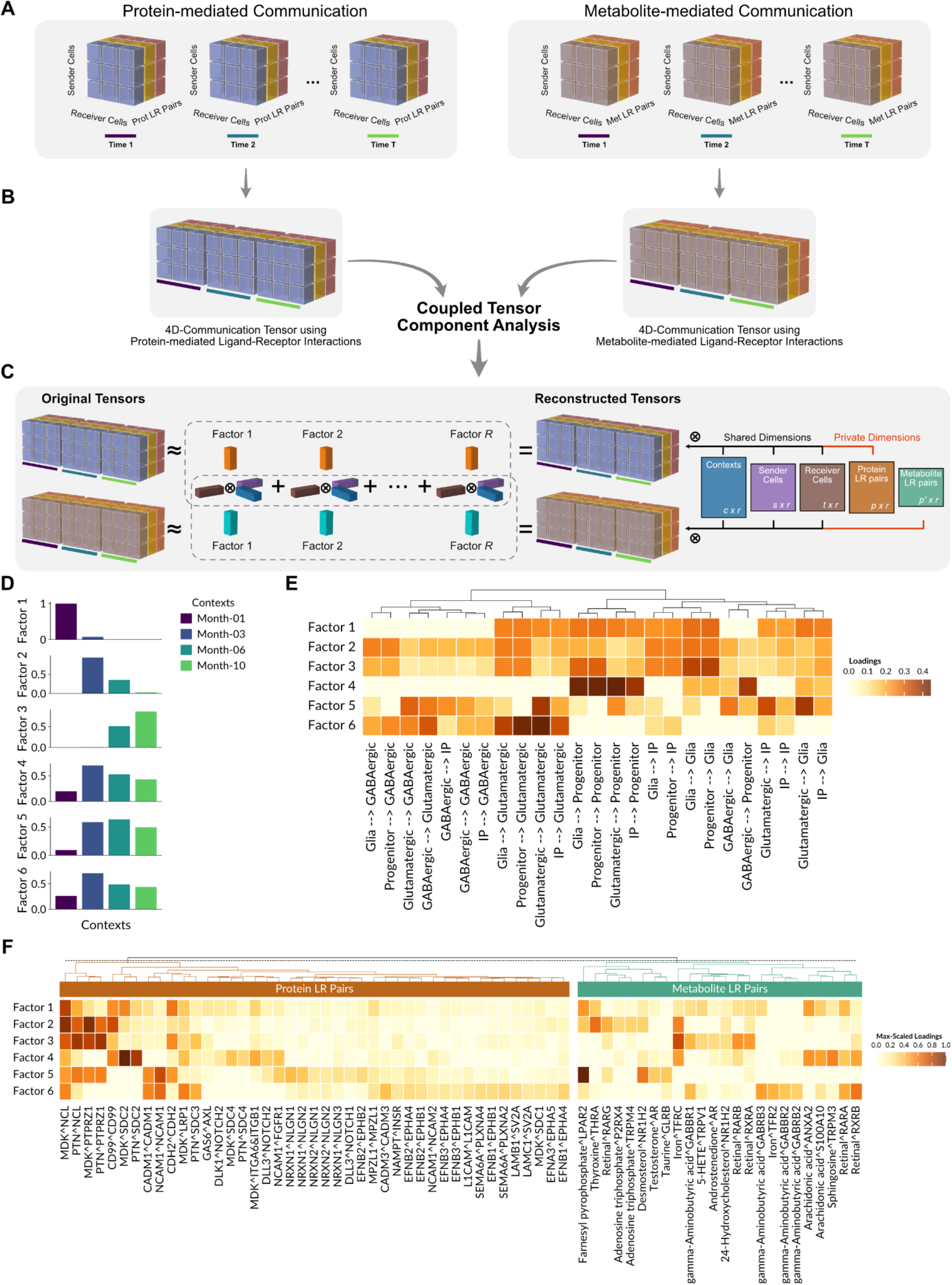
CTCA for detecting coordinated protein- and metabolite-based communication patterns. (**A**) 4D tensors are built separately for protein- and metabolite-mediated communication modalities by computing communication scores for each ligand-receptor (LR) pair in every combination of sender-receiver cell pairs at each time point, using cell2cell and MEBOCOST, respectively. (**B**) These tensors are jointly analyzed with our non-negative coupled tensor component analysis (CTCA). (**C**) CTCA decomposes both tensors into R factors, each of which is the outer product (⊗) among loadings vectors (colored boxes) of the shared and private tensor dimensions. Shared dimensions are contexts, sender cells, and receiver cells, and their loading vectors (blue, purple, and brown boxes), couple both modalities, while loading vectors for private dimensions (orange and cyan boxes) capture protein and metabolite-specific LR pairs. Summing these factors reconstructs each tensor, which can also be represented as outer products of matrices (colored rectangles), each specific to one dimension, containing loadings for the dimension elements (rows) across factors (columns), as indicated by their size at their bottom right corner. (**D**–**F**) Application of CTCA to cortical brain organoid scRNA-seq data reveals six CCC programs (factors). (**D**) Context loadings per factors reflect shared dynamics of CCC between modalities. (**E**) Heatmap of sender-receiver cell pair loadings per factor, hierarchically clustered. These joint loadings were computed as the outer product of the shared loadings of sender and receiver cell dimensions. (**F**) Heatmaps of protein- and metabolite-specific LR loadings per factor, hierarchically clustered separately by ligand type. Only LR pairs with high loadings in at least one factor are displayed. Co-occurrence of high-loading protein and metabolite LR pairs within the same factor reflects their coordinated communication dynamics. Loadings were scaled to the maximum within each modality. IP, intermediate progenitor; LR, ligand-receptor.

To illustrate the utility of the CTCA, we applied it on a dataset of cortical brain organoids containing multiple time points that represent different stages of cortical brain development (Trujillo *et al*., 2019). This complex temporal process involves a sequence of stages where neural and non- neural cell types must be produced in the correct number, and present proper spatiotemporal dynamics of CCC (Jiang and Nardelli, 2016). After building both coupled tensors for this dataset (Figures 1A-C), we used Tensor-cell2cell v2 to detect six CCC programs coordinating both ligand types (factors in Figures 1D-F). These programs capture distinct dynamics during the development of cortical brain organoids, involving either specific time points (factors 1-3) or changes across time points (factors 4-6) (Figure 1D). Factor 1 was dominated by the first month, factor 2 peaked at month 3 and declined by month 6, while factor 3 emerged at month 6 and peaked later. Factors 4-6 are examples of communication programs present across time, with a peak in the mid stages.

The CTCA is key to identify coordination from two modalities across specific temporal dynamics and sender-receiver cells. Among factors with prominent and specific CCC patterns, factor 4 was dominated by communication from different sender cells to progenitors and to a lesser extent to glial cells as receivers (Figure 1E), reflecting a progenitor-centered communication hub. Indeed, it was characterized by progenitor-supporting signals, including CD99–CD99 self-signaling, MDK–SDC2 and PTN–SDC2 growth factor interactions, and DLL3/DLK1–NOTCH2 in the protein modality, acting in concert with metabolite LR pairs such as iron–TFRC, arachidonic acid– ANXA2/S100A10, sphingosine–TRPM3, and retinal–RARA/RXRB (Figure 1F), consistent with programs that sustain and regulate progenitor populations (Zou *et al*., 2006; Borghese *et al*., 2010; Sakayori *et al*., 2011). In contrast, factor 6 was defined by signaling into glutamatergic and GABAergic neurons from multiple sender types (Figure 1E), indicating a neuronal-centered program rather than progenitor-focused communication. This CCC program was characterized by neuronal adhesion and synaptic ligand-receptor pairs (e.g., Laminins and Ephrins interactions with their cognate receptors, NCAM1 and CADM1 self-interactions, and Neurexin–Neuroligin interactions) together with metabolites such as retinoids and GABA (Figure 1E), providing mechanistic detail for the neuronal-centered program. Thus, our results highlight how CTCA reveals the coordinated action of diverse molecular signals that guide progenitor maintenance and neuronal maturation.

## 4 Discussion

We demonstrate through our implementation of a non-negative CTCA that Tensor-cell2cell v2 can decipher the coordinated temporal dynamics of both protein- and metabolite-mediated CCC from single-cell omics. Unlike existing methods, it jointly analyzes modalities with distinct behaviors, such as protein-based communication (directly predicted from gene expression) and metabolite-based communication (indirectly inferred from enzyme expression). Tensor-cell2cell builds modality-specific 4D communication tensors and couples them through CTCA, aligning their dynamics across contexts and sender-receiver cell pairs, which would otherwise show mismatched behaviors if analyzed separately. This facilitates the identification of key LR pairs across ligand types acting in concert within pertinent CCC programs.

Applied to cortical brain organoid development (Trujillo *et al*., 2019), our approach revealed concerted dynamics of specific protein and metabolite ligands. For example, both MDK and retinal interactions with their cognate receptors were repeatedly captured across factors (Figure 1F). MDK is encoded by a retinoic acid (RA)-responsive gene (Kadomatsu and Muramatsu, 2004), and retinal is the precursor of RA, a key developmental regulator (Maden, 2007). CTCA also identified progenitor-centered programs involving Notch and arachidonic acid signaling (factor 4 in Figure 1F), whose coordinated action help maintaining a proliferative state instead of undergoing differentiation (Sakayori *et al*., 2011; Greig *et al*., 2013; Ohtaka-Maruyama and Okado, 2015; Jiang and Nardelli, 2016). These findings illustrate how Tensor-cell2cell v2 uncovers biologically meaningful coordination between protein- and metabolite-mediated communication.

Altogether, Tensor-cell2cell v2 provides a flexible framework to integrate multiple CCC modalities. Beyond proteins and metabolites, CTCA can also couple other diverse modalities, such as extracellular vesicle signaling (Shao *et al*., 2025), transcription factor activities (Badia-I-Mompel *et al*., 2022), or intracellular metabolism (Armingol *et al*., 2025), with classical communication scores (Armingol, Baghdassarian, *et al*., 2022). Although we used cell2cell (Armingol, Ghaddar, *et al*., 2022) and MEBOCOST (Zheng *et al*., 2025), users can adopt alternative tools for each modality, depending on their needs and each tool’s underlying assumptions. By harmonizing heterogeneous scores into multimodal CCC programs, Tensor-cell2cell v2 facilitates the discovery of coordinated molecular mediators shaping cellular contexts, and enables future multimodal analyses.

## Acknowledgements

This work was supported by NIGMS grant R35 GM119850 to NEL, and by the NVIDIA Corporation through its Academic Hardware Grant Program to EA. EA was also supported by the Chilean Agencia Nacional de Investigación y Desarrollo (ANID) through its scholarship program DOCTORADO BECAS CHILE/2018 - 72190270, the Fulbright Chile Commission, and the Siebel Scholar Foundation. ROL was supported in part by the UCSD Graduate Training Program in Cellular and Molecular Pharmacology through an institutional training grant from the National Institute of General Medical Sciences, T32 GM007752. HMB was supported in part by an appointment to the Research Participation Program at the U.S. Food and Drug Administration administered by the Oak Ridge Institute for Science and Education through an interagency agreement between the U.S. Department of Energy and the U.S. Food and Drug Administration.

