## Supplementary Figure 1 for "Tensor-cell2cell v2 unravels coordinated dynamics of protein- and metabolite-mediated cell-cell communication"

### Content:

- Supplementary Figure S1

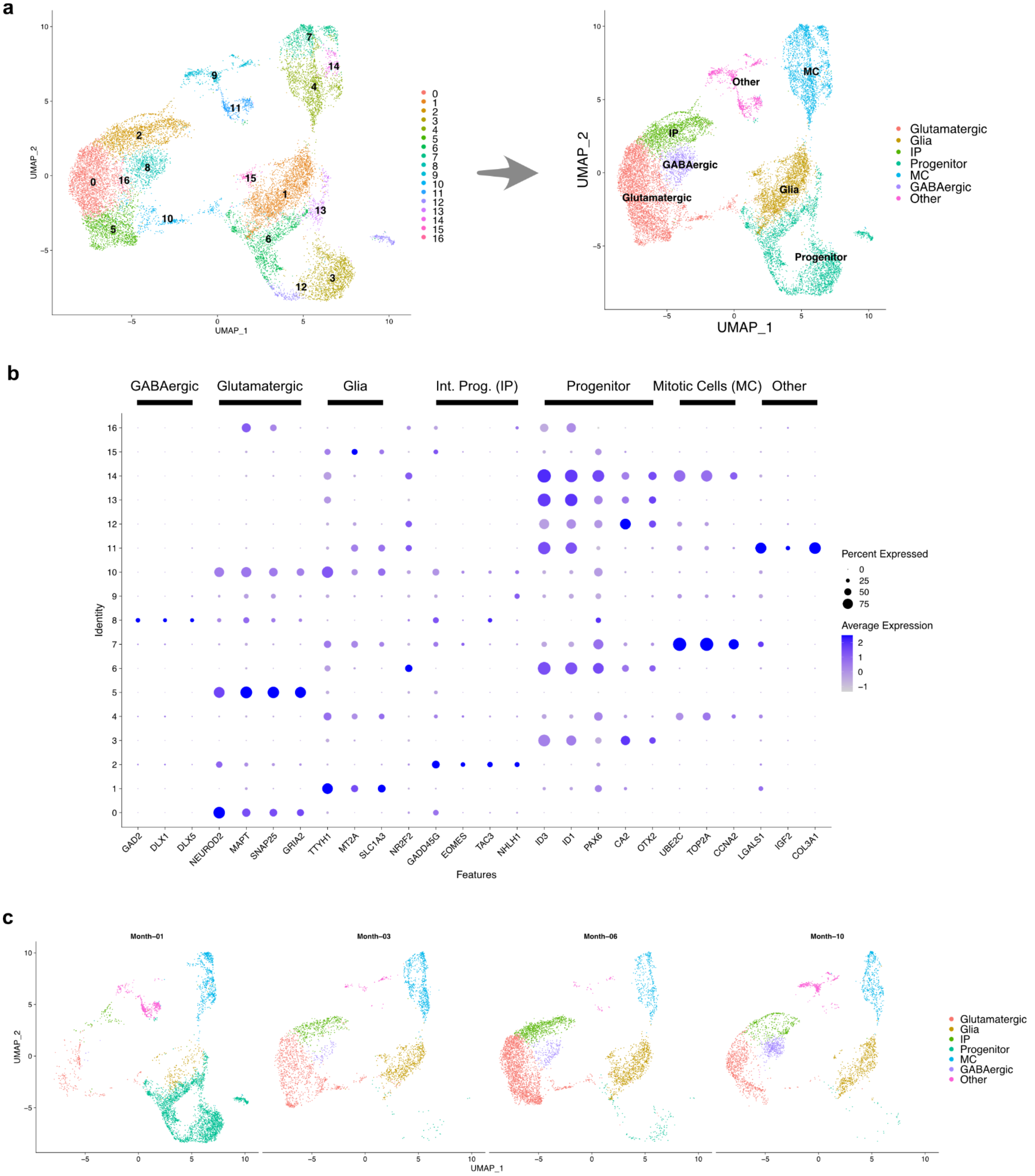


**Supplementary Figure S1. Annotation of the single-cell dataset of the brain cortical organoid.** After preprocessing the dataset and performing the cell clustering as indicated in Methods, each of the clusters was annotated. (**A**) UMAP visualization of the resulting clusters and their annotations. (**B**) Detail of the marker expression levels employed to perform the annotation. These markers were previously described by (Trujillo et al., 2019). (**C**) UMAP visualization of the distinct cell types across the different time points in the dataset.
